# PRMT7 ablation stimulates anti-tumor immunity and sensitizes melanoma to immune checkpoint blockade

**DOI:** 10.1101/2021.07.28.454202

**Authors:** Nivine Srour, Oscar D. Villarreal, Zhenbao Yu, Samuel Preston, Wilson H. Miller, Magdelena M. Szewczyk, Dalia Barsyte-Lovejoy, Han Xu, Sonia V. del Rincón, Stéphane Richard

## Abstract

Despite the success of immune checkpoint inhibitor (ICI) therapy in different cancers, resistance and relapses are frequent. Thus, combination therapies are expected to enhance response rates and overcome resistance to ICIs. Herein, we report that combining protein arginine methyltransferase 7 (PRMT7) inhibition with ICIs triggers a strong anti-tumor T cell immunity and restrains tumor growth *in vivo* by increasing tumor immune cell infiltration. Consistently, TCGA database analysis showed an inverse correlation between PRMT7 expression and T cell infiltration in human melanomas. Mechanistically, we show that PRMT7 has a two-prong effect on melanoma tumor immunity. On one hand, it serves as a coactivator of IRF-1 for PD*-L1* expression by upregulating promoter H4R3me2s levels in melanoma cells. Next, PRMT7 prevents repetitive element expression to avoid intracellular dsRNA accumulation or ‘viral mimicry’. PRMT7 deletion resulted in increased endogenous retroviral elements (ERVs), dsRNA, and genes implicated in interferon activation, antigen presentation and chemokine signaling. Our findings identify PRMT7 as factor used by melanoma to evade anti-tumor immunity and define the therapeutic potential of PRMT7 alone or in combination with PD-(L)1 blockade to enhance ICI efficiency.

## Introduction

Arginine methylation is an epigenetic modification dysregulated in cancer with the frequent overexpression of protein arginine methyltransferases (PRMTs) (Yang and Bedford, 2013). There are three type of PRMTs, type I, II, and III, generating as final products asymmetric dimethylarginine (ADMA), symmetric dimethylarginine (SDMA) and monomethylarginine (MMA), respectively (Bedford and Clarke, 2009). High affinity specific PRMT inhibitors have been developed and are in clinical trials (Guccione and Richard, 2019). The goal is to understand which cancer patients would best benefit from these PRMT inhibitors either alone or in combination therapies (Wu et al., 2021). Links between PRMTs and immune development and function have been identified, however, the role of arginine methylation in immunotherapy is emerging.

Arginine methylation by PRMT1 and PRMT5 have been shown to regulate immune function. PRMT1 functions as a coactivator of RORγt for Th17 differentiation (Sen et al., 2018) and interacts with GFI1, a transcriptional regulator required for development and maintenance of T lymphoid leukemia to regulate the DNA damage response (Vadnais et al., 2018). Deletion of PRMT1 in mature B cells results in reduced activation and differentiation, and impairs humoral immunity (Infantino et al., 2017). Furthermore, PRMT1 regulates pre-B cell differentiation via the methylation of cyclin-dependent kinase 4 (CDK4) (Dolezal et al., 2017). PRMT5 modulates T cell activation processes via the regulation of the transcription of cytokine genes induced by interferon (IFN) (Metz et al., 2020), and when deleted, PRMT5 decreases signaling via γc-family cytokines and reduces peripheral CD4^+^ T cells and CD8^+^ T cells (Inoue et al., 2018). PRMT5 inhibition also blunts adaptive memory Th cell responses and reduces inflammation in the EAE (Experimental Autoimmune Encephalomyelitis) mouse model (Webb et al., 2017). PRMT5 regulates B cells via the regulation of the germinal center reaction and the antibody response (Litzler et al., 2019). Recently, PRMT5 was shown to methylate cGAS, which in turn abolish its DNA binding ability and attenuates the antiviral immune response (Kim et al., 2020; Ma et al., 2021). Taken together, inhibition of arginine methylation is an attractive therapeutic approach for B and T cell-mediated disease (Parry and Ward, 2010).

The epigenetic modifier PRMT7 catalyzes MMA mainly on histones proteins (Zurita-Lopez et al., 2012; Feng et al., 2013a; Jain and Clarke, 2019) and the methylation of histone H4 by PRMT7 was shown to allosterically modulate the ability of PRMT5 to generate H4R3me2s (Feng et al., 2013a; Jain et al., 2017). Genetic loss-of-function *PRMT7* mutations and deletions cause the SBIDDS (short stature, brachydactyly, intellectual developmental disability and seizures syndrome) syndrome (Agolini et al., 2018). PRMT7-null mice have impaired muscle (Blanc et al., 2016; Jeong et al., 2016), adipogenesis (Leem et al., 2019) and B cell germinal center formation (Ying et al., 2015). Zebrafish PRMT7 was shown to negatively regulate the antiviral response (Zhu et al., 2020), linking arginine methylation to the modulation of innate and adaptive immunity. Little is known about the role of PRMT7 in immune regulation, herein, we identify PRMT7 as a modulator of immunotherapy in melanoma.

Tumor cells evade antitumoral immunosurveillance and this has led to development of immune checkpoint inhibitors (ICIs). One prominent axis by which this occurs is the blockade of programmed death-protein 1 (PD-1) and its ligand (PD-L1) (Wherry and Kurachi, 2015). PD-L1, known also as *CD274* and *B7-H1* (Dong et al., 1999), is a transmembrane protein expressed on cancer cells that binds PD-1 on B cells, T cells and myeloid cells (Dong et al., 2002). Therapeutic approaches targeting PD-1, PD-L1 (Pardoll, 2012; Salmaninejad et al., 2019) and the cytotoxic T-lymphocyte-associated antigen 4 (CTLA-4) have proven effective towards activating the host immune system in many cancers including melanoma (Larkin et al., 2015; Topalian et al., 2016; Larkin et al., 2019). However, approximately 50% of patients fail to respond or acquire resistance to ICI therapy (Feng et al., 2013b; Chen and Mellman, 2017). The combination of ICI and epigenetic inhibitors holds promise to fill in this therapeutic gap. Inhibitors such as DNA methylation inhibitors 5-azacytidine (Aza), 5-aza-2′-deoxycytidine (5-Aza-dC; Decitabine) (Bian and Murad, 2014; Chiappinelli et al., 2015) or the histone lysine demethylase LSD1 inhibition (Sheng et al., 2018) are known to increase immune signaling (e.g. activation of IFN pathway and secretion of cytokines).

Elevated PD-L1 expression on cancer cells provides evasion from T cell controlled immune surveillance. Herein, we provide evidence that PRMT7 functions as a dual regulator, 1) an epigenetic coactivator of IRF-1 for *PD-L1* expression by upregulating promoter H4R3me2s levels in melanoma, and 2) PRMT7 plays a role in suppressing the expression of endogenous repetitive sequences to maintain low intracellular dsRNA levels to prevent an anti-viral response. Both these tumor intrinsic PRMT7 functions are complementary to enhance immune evasion and affect the sensitivity to ICI therapy. In sum, PRMT7 deletion or inhibition leads to potent anti-tumor T cell immunity and renders melanomas more responsive to anti-CTLA-4 and anti-PD-1 therapy.

## Results

### PRMT7 deficiency enhances sensitivity to anti–CTLA-4 and anti-PD-1 therapy *in vivo*

Previously, a CRISPR/Cas9 genetic screen in a murine model of melanoma (B16.F10) was performed to identify genes that, when deleted, improve anti-tumor responses to immunotherapy (Manguso et al., 2017). For this, tumor cells were injected into mice that were treated with GVAX (granulocyte macrophage colony-stimulating factor (GM-CSF)-secreting, irradiated tumor cell vaccine) alone or in combination with monoclonal PD-1 blockade to improve the immune system by stimulating anti-tumor T cell infiltration and myeloid cell activation. Interestingly, PRMT7 ranked 8^th^ in the GVAX + PD-1 vs TCRα*^−/−^* CRISPR/Cas9 screen and ranked 23^rd^ in the GVAX vs TCRα*^−/−^* CRISPR/Cas9 screen (Manguso et al., 2017). We re-analyzed their CRISPR/Cas9 screen data using our developed software called MoPAC (Gao et al., 2019), and PRMT7 remained a top hit (Supplementary Fig. S1A, S1B and Dataset S1). Moreover, we identified elevated *PRMT7* mRNA expression in many types of cancers and high *PRMT7* levels were associated with reduced patient survival for melanoma (Supplementary Fig. S2A-C). Thus, we hypothesized that the PRMT7 epigenetic regulator promotes immunosuppression in melanoma.

To examine whether PRMT7 deficiency enhanced susceptibly to CTLA-4 and PD-1 blockade in melanoma, we generated two CRISPR/Cas9 PRMT7 depleted B16.F10 melanoma clones (sgPRMT7-1, sgPRMT7-2). The lack of PRMT7 was shown by immunoblotting and RT-qPCR (Fig. 1A, 1B) and the PRMT7 depleted cell lines displayed similar growth rates *in vitro* as control (sgCTL) cells (Supplementary Fig. S3). We implanted sgCTL and PRMT7 depleted B16.F10 cells (sgPRMT7-1, sgPRMT7-2) subcutaneously in syngeneic C57BL6/J mice and monitored tumor growth in the absence or presence of CTLA-4 and PD-1 monoclonal antibodies (Fig. 1C-E), a combination more effective than monotherapies (Wei et al., 2019). Without ICI treatment, we observed a small but significant difference in tumor initiation between the sgCTL and sgPRMT7 melanoma tumors at days 3, 6 and 9, but not at the day 15 endpoint (Fig. 1C). However, treatment with CTLA-4 and PD-1 monoclonal antibodies showed a markedly reduced tumor size (> 90% at day 18) in both sgPRMT7 cell lines compared to the tumors generated by sgCTL cells (Fig. 1D). Similar tumor growth was observed at day 15 in TCRα^-/-^ mice (Fig. 1E) without CTLA-4 and PD-1 blockade, suggesting that sgPRMT7 cells elicited a potent anti-tumor T cell immunity *in vivo*, rather than affecting tumor cell growth. These data demonstrate a synergy between PRMT7 deletion and immune checkpoint inhibitors in controlling melanoma tumor growth.

**Figure 1:**
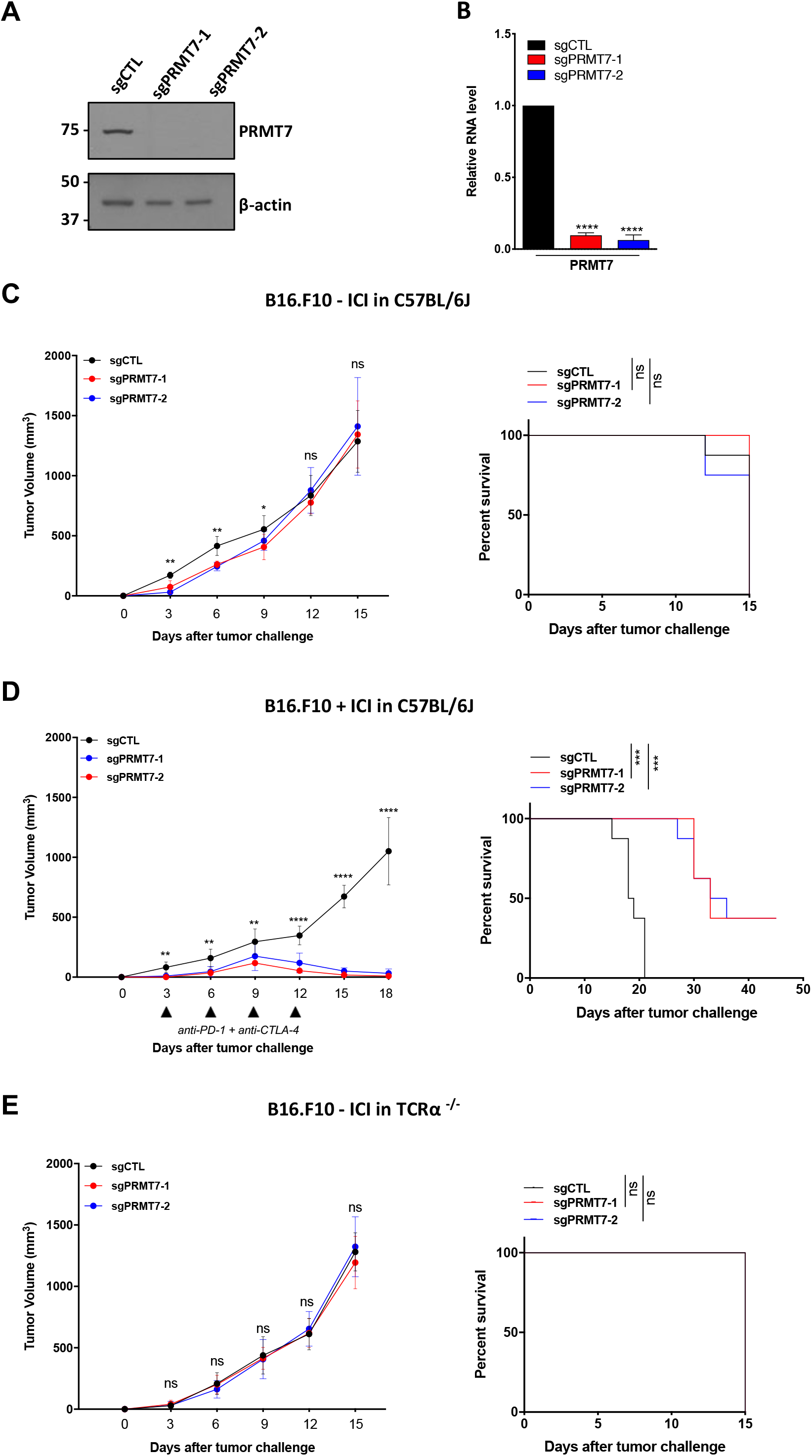
Deletion of PRMT7 sensitizes B16.F10 melanomas to ICI. **(A)** Western blot showing the expression of PRMT7 in sgCTL and sgPRMT7-1 and 2 targeted B16.F10 melanoma cells. β-actin is the loading control. Molecular mass markers are indicated in the left in kDa. One representative image out of three is shown. **(B)** PRMT7 mRNA levels for clones in **(A)** measured by RT-qPCR. Data are mean ± SD. Data representative for four independent experiments. Statistical significance was calculated by unpaired student *t* test (\*\*\*\**p* <0.0001). **(C)** Left panel: Tumor volume averaged for each group at each time point for sgCTL, sgPRMT7-1 and sgPRMT7-2 B16 cells injected into C57BL/6J mice without ICI treatment. Right panel: Kaplan-Meier survival curves were assessed at indicated time points. All groups reached the endpoint on the same day. Data are mean ± SEM; *n* = 8-10 mice per group.; *p* values were determined using multiple *t* test (* *p* <0.05; \*\**p* <0.01; *ns*: non-significant). **(D)** Left panel: Tumor volume averaged for each group at each time point for sgCTL, sgPRMT7-1 and sgPRMT7-2 B16 cells injected into C57BL/6J mice treated intraperitoneally with anti-CTLA-4 and anti-PD-1 (ICI) at day 3, 6, 9 and 12 (black triangles). Right panel: Kaplan-Meier survival curve was assessed at indicated time points. Data are mean ± SEM; *n* = 8-10 mice per group. Representative of two to three independent experiments is shown; *p* values were determined using multiple *t* test (\*\**p* <0.01; \*\*\**p* <0.001; \*\*\*\**p* <0.0001). **(E)** Left panel: Tumor volume averaged for each group at each time point for sgCTL, sgPRMT7-1 and sgPRMT7-2 B16 cells injected into TCRα KO mice without ICI treatment. Right panel: Kaplan-Meier survival curves were assessed at indicated time points. All groups reached the endpoint on the same day. Data are mean ± SEM; *n* = 8-10 mice per group.; *p* values were determined using multiple *t* test (*ns*: non-significant).

We next tested the PRMT7 inhibitor, SGC3027, a cell active prodrug, which in cells is converted to the active compound SGC8158 (Szewczyk et al., 2020). Since the prodrug is expected to have poor systemic pharmacokinetic properties, for the proof-of-concept we opted for intra-tumoral delivery. B16.F10 melanoma were injected subcutaneously in C57BL/6J mice and on day 7, mice were treated with 10 µM of DMSO, SGC3027N (inactive compound) or SGC3027 (active compound) via intratumoral injection (4 doses for 4 days). Interestingly, we found that 96 hours after the last injection with SGC3027, the tumor growth significantly decreased as compared to mice injected with DMSO alone or with SGC3027N (Supplementary Fig. S4). These data suggest that PRMT7 inhibition is of therapeutic value to potentiate the effect of ICI therapy.

### Intrinsic loss of PRMT7 in tumors promotes T cell infiltration and regulates melanoma cell plasticity

To assess whether the difference in tumor growth observed in sgPRMT7 melanomas was due to differential immune cell infiltration into tumors, we evaluated the immune composition of sgCTL and sgPRMT7 melanomas following anti-CTLA-4 and -PD-1 treatment. We focused on myeloid derived suppressor cell populations (MDSCs), which are known to mediate resistance in ICI therapy (Hou et al., 2020). Using multi-parameter flow cytometry (Supplementary Fig. S5), we observed a lower level of recruitment of granulocytic (G)-MDSCs and monocytic (M)-MDSCs in sgPRMT7 melanomas (Fig. 2A, 2B) with no difference in the total number of CD3^+^ T cells or non-lymphatic dendritic cells (NLT DCs, Fig. 2C, 2D). Moreover, we observed an elevated number of infiltrating CD3^+^CD8^+^ T cells in tumors from mice implanted with sgPRMT7 treated with ICI therapy (Fig. 2E-H). Together the decrease in MDSCs and the increase in CD8^+^ T cells play a role in reducing tumor growth of sgPRMT7 B16.F10 cells and markedly enhance adaptive immunity following anti-CTLA-4 and PD-1 treatment.

**Figure 2:**
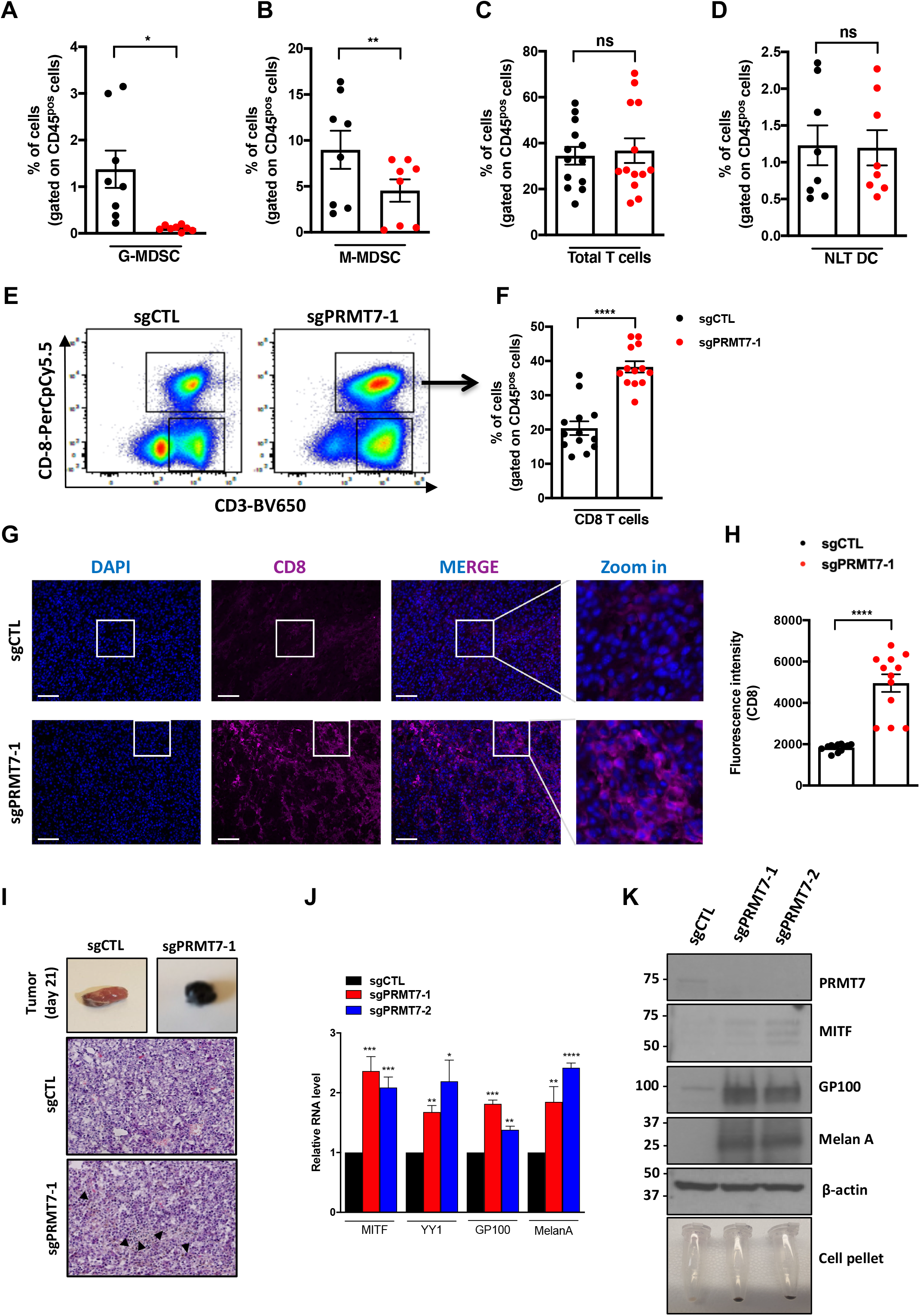
Deletion of PRMT7 in melanomas increases immune cell infiltration and increases melanocytic plasticity. **(A-D)** Tumors were digested into a single cell suspension and their immune cell composition analyzed. Quantification of Myeloid-Derived Suppressor Cell (MDSC) populations such as Granulocytic MDSC (G-MDSC, F4/80^neg^) **(A)**, Monocytic MDSC (M-MDSC, F4/80^pos^) **(B)**, total T cells **(C)** and non-lymphatic dendritic cells (NLT DC) **(D)** in sgCTL (black) and sgPRMT7-1 (red) B16.F10 tumors. **(A-D)** The data represents the mean ± SD and is from two to three independent experiments with a minimum of 3 mice per group. Each dot represents one mouse. Statistical significance was calculated using paired student *t* test. (* *p* <0.05; \*\**p* <0.01; *ns*: non-significant). **(E)** Representative flow cytometry plots using anti-CD45, anti-CD3 and anti-CD8 antibodies in sgCTL and sgPRMT7-1 B16.F10 tumors. **(F)** Quantification graphs from (**E)** showing frequencies of double positive CD3^pos^, CD8^pos^ cells (gated on CD45^pos^ cells) in sgCTL (black) and sgPRMT7-1 (red) B16.F10 tumors. Cells were gated as indicated and the relative percentage of cells shown; *n*=13 animals per group; data from three independent experiments. Statistical significance was calculated using paired student *t* test (\*\*\*\**p* <0.0001). **(G)** Representative immunofluorescent images of sgCTL and sgPRMT7-1 tumor sections treated with anti-CTLA-4 and anti-PD-1 *in vivo* and stained with anti-CD8α antibody at 10x magnification. DAPI, 4’,6-diamidino-2-phenylindole, was used to visualize nuclei by Zeiss confocal microscopy. A minimum of 3 biological replicates were used for each experiment. **(H)** Immunofluorescence intensity of the CD8 staining done in (**G**) using ImageJ software. Bar graphs show fluorescence mean intensity ± SEM. Statistical significance was calculated using unpaired student *t* test (\*\*\*\**p* <0.0001). **(I)** Representative pictures of subcutaneous sgCTL and sgPRMT7-1 derived-melanomas in C57BL/6J mice treated with anti-CTLA-4 and anti-PD-1 (day 21, top) and corresponding representative images of H&E-stained tumor sections (day 21, bottom). Black arrowheads indicate the pigmented areas. **(J)** RT-qPCR analysis of *Mitf*, *Yy1*, *Gp100* and *Melan-A* mRNA transcripts in sgCTL, sgPRMT7-1 and sgPRMT7-2 B16.F10 cells. Data are mean ± SD. Data representative for 3 independent experiments. Statistical significance was calculated by unpaired student *t* test (* *p* <0.05; \*\**p* <0.01; \*\*\**p* <0.001; \*\*\*\**p* <0.0001). **(K)** Western blot showing the expression of the indicated proteins (PRMT7, MITF, GP100 and Melan-A) in sgCTL, sgPRMT7-1 and sgPRMT7-2 B16.F10 melanoma cells. β-actin was used as the loading control. Molecular mass markers are indicated in the left in kDa. Data are representative of three independent experiments. Cell pellet representative images were shown (bottom; note the black pellet in sgPRMT7 clones).

Melanomas with high immune infiltrates have been associated with a pigmented, differentiated phenotype (Wiedemann et al., 2019). Pigmentation is regulated by MITF (microphthalmia-associated transcription factor), a determinant of melanoma cell plasticity (Du et al., 2003). Dedifferentiated melanomas, characterized by low MITF expression are generally invasive and resistant to immunotherapy (Hoek et al., 2006; Hoek and Goding, 2010). Since PRMT7 null tumors exhibited an increase in immune cell infiltration and were sensitized to immunotherapy, we postulated that PRMT7 might influence plasticity of melanoma cells (Fig. 2I-K). The sgPRMT7 derived tumors were indeed more pigmented, compared to sgCTL tumors (Fig. 2I). sgPRMT7 tumors showed an increase in MITF mRNA and protein levels and two other melanocytic antigens, Melan-A (also known as MART-1) and GP100 (also known as Pmel17, Fig. 2J, 2K). These findings suggest that PRMT7 regulates melanoma cell plasticity by modulating melanocyte antigen gene expression.

### PRMT7 loss represses PD-L1 expression in melanoma

The decreased tumor size in mice implanted with sgPRMT7 B16.F10 cells and receiving ICI therapy (Fig. 1D) suggested that PRMT7 may regulate the PD-1 axis. To examine whether PRMT7 affected PD-L1 expression, B16 melanoma cells were transfected with control siRNA against an irrelevant gene, Firefly luciferase, (siLuc) or a Smartpool of PRMT7 siRNAs. A decrease in PD-L1 protein expression level was observed in siPRMT7 compared to siLuc transfected B16.F10 melanoma cells (Fig. 3A). This decrease was also observed at the cell surface of siPRMT7 cells using flow cytometry (Fig. 3B). The decrease in *PD-L1* mRNA was observed in siPRMT7 transfected cells or treatment with SGC3027 (Fig. 3C, 3D). In contrast, siPRMT5 transfected cells (Fig. 3C) or treatment with the PRMT5i EPZ015666 (Fig. 3D) increased the expression of *PD-L1* mRNA, as previously reported (Kim et al., 2020). In addition, we tested other PRMT inhibitors, but we did not observe any significant differences in *PD-L1* mRNA expression with a type I PRMTi (MS023) and CARM1i (TP064, Fig. 3D). We also confirmed the reduced *PD-L1* mRNA and protein levels in sgPRMT7-1 and sgPRMT7-2 (Fig. 3E, 3F). The ectopic expression of GFP-PRMT7 in sgPRMT7-1 partially rescued the *PD-L1* mRNA and protein levels (Fig. 3G, 3H).

**Figure 3:**
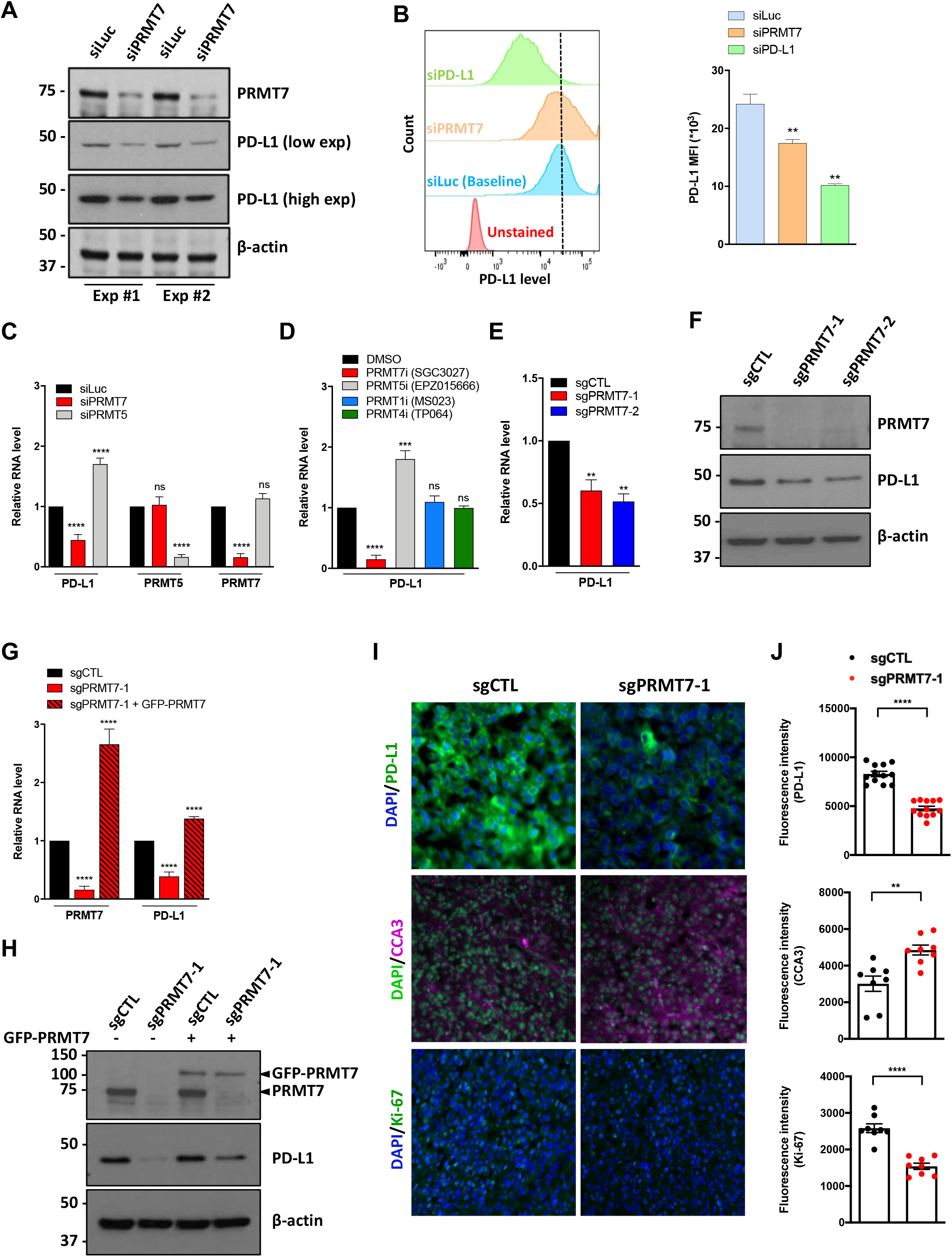
PRMT7 positively regulates PD-L1 expression in melanoma. **(A)** Western blot analysis of PRMT7 and PD-L1 expression in total cell lysates isolated from B16.F10 cells transfected with either siRNA targeting luciferase (siLuc) or PRMT7 (siPRMT7). β-actin was used as the loading control. Two independent experiments are shown (Exp #1, Exp #2). Two exposures of PD-L1 expression are shown (low and high exposure). Molecular mass markers are indicated in the left in kDa. **(B)** Left panel: Flow cytometry histograms showing PD-L1 surface expression at baseline level (blue) and in B16.F10 cells transfected with either siPRMT7 (orange) or siPD-L1 (positive control; green). Right panel: The mean fluorescence intensity (MFI) of PD-L1^pos^ cells by flow cytometry is shown. Statistical significance was calculated by unpaired student *t* test (*\*\*p* <0.01). **(C)** *PD-L1*, *PRMT5* and *PRMT7* mRNA abundance measured by RT-qPCR in siLuc (black), siPRMT5 (grey) and siPRMT7 (red) transfected B16.F10 cells is shown. Bar graphs represent the mean fold-change ± SD. Data are representative of three independent experiments. Statistical significance was calculated by unpaired student *t* test (\*\*\*\**p* <0.0001; *ns*: non-significant). **(D)** *PD-L1* mRNA abundance measured by RT-qPCR in melanoma cells treated with the indicated PRMT inhibitors for 48h (SGC3027: 10 µM; EPZ015666: 5 µM; MS023: 600 nM and TP064: 3 µM). Bar graphs represent the mean fold-change ± SD. Data are representative of three independent experiments. Statistical significance was calculated by unpaired student *t* test (\*\*\**p* <0.001; \*\*\*\**p* <0.0001; *ns*: non-significant). **(E)** *PD-L1* mRNA abundance measured by RT-qPCR in sgCTL, sgPRMT7-1 and sgPRMT7-2 melanoma cells. Bar graphs represent the mean fold-change ± SD. Data are representative of three independent experiments. Statistical significance was calculated by unpaired student *t* test (\*\**p* <0.01). **(F)** Western blot analysis of PRMT7 and PD-L1 expression in total cell lysates isolated from sgCTL, sgPRMT7-1 and sgPRMT7-2 cells. β-actin was used as the loading control. The data is representative of greater than three independent experiments. Molecular mass markers are indicated in the left in kDa. **(G)** RT-qPCR analysis of *PRMT7* and *PD-L1* mRNAs in sgCTL, sgPRMT7-1 and sgPRMT7-1 + GFP-PRMT7 (rescued) cells. Data are representative for three independent experiments. **(H)** Western blot analysis of PRMT7 and PD-L1 expression in total cell lysates isolated from sgCTL, sgPRMT7-1, sgCTL + GFP-PRMT7, and sgPRMT7-1 + GFP-PRMT7 cells. β-actin was used as the loading control. A representative experiment is shown, and the experiment was performed twice. The molecular mass markers are indicated in the left in kDa. **(I)** PRMT7-deficient melanomas show less PD-L1 expression *in vivo*, less ki-67and more cleaved-caspase-3 (CCA3). Representative immunofluorescent images showing the expression of PD-L1 (20x), CCA3 and ki-67 at 40x magnification in sgCTL and sgPRMT7-1 derived melanoma tumor sections from mice treated with immunotherapy drugs *in vivo*. DAPI, 4’,6-diamidino-2-phenylindole, was shown in blue or green as indicated. **(J)** Immunofluorescence intensity of the staining performed in (**I**) using ImageJ software. Bar graphs show mean intensity ± SEM. Statistical significance was calculated by unpaired student *t* test (\*\**p* <0.01; \*\*\*\**p* <0.0001).

Moreover, immunostaining using anti-PD-L1 antibody on sgCTL and sgPRMT7-1 tumor sections further confirmed the reduced PD-L1 expression (Fig. 3I, 3J). This correlated with lower proliferation, as visualized by reduced Ki-67 staining, and higher level of apoptosis, as an increase in cleaved caspase 3 (CCA3) in PRMT7-deficient tumors *in vivo* was observed (Fig. 3I, 3J). These findings define PRMT7 as a positive regulator of *PD-L1* expression and show that this property is unique to PRMT7.

### PRMT7 functions as a coactivator of IRF-1 on the *PD-L1* promoter

We tested whether PRMT7 localized at the *PD-L1* gene using chromatin immunoprecipitation (ChIP) assay in B16.F10 cells. We used five known *PD-L1* regulatory regions R1-R5 spanning the *PD-L1* promoter from −4,000 to +1,000 Kb (Fig. 4A) and we showed that PRMT7 bound distal regions R1, R2 and R3, but not R4 and R5 of the *PD-L1* promoter (Fig. 4B). We next performed ChIP on IRF-1, a transcription factor responsible for IFN-γ-induced *PD-L1* expression (Lu et al., 2016). IRF-1 bound to the *PD-L1* regions R1-R3, but not R4 and R5, with no difference at the *β-actin* promoter, as control (Fig. 4C). Interestingly, siPRMT7 B16.F10 melanoma cells displayed reduced IRF-1 binding at the *PD-L1* promoter R1-R3 regions (Fig. 4C). These findings suggest the presence of PRMT7 at the *PD-L1* promoter R1-R3 regions influences IRF-1 binding and/or recruitment.

**Figure 4:**
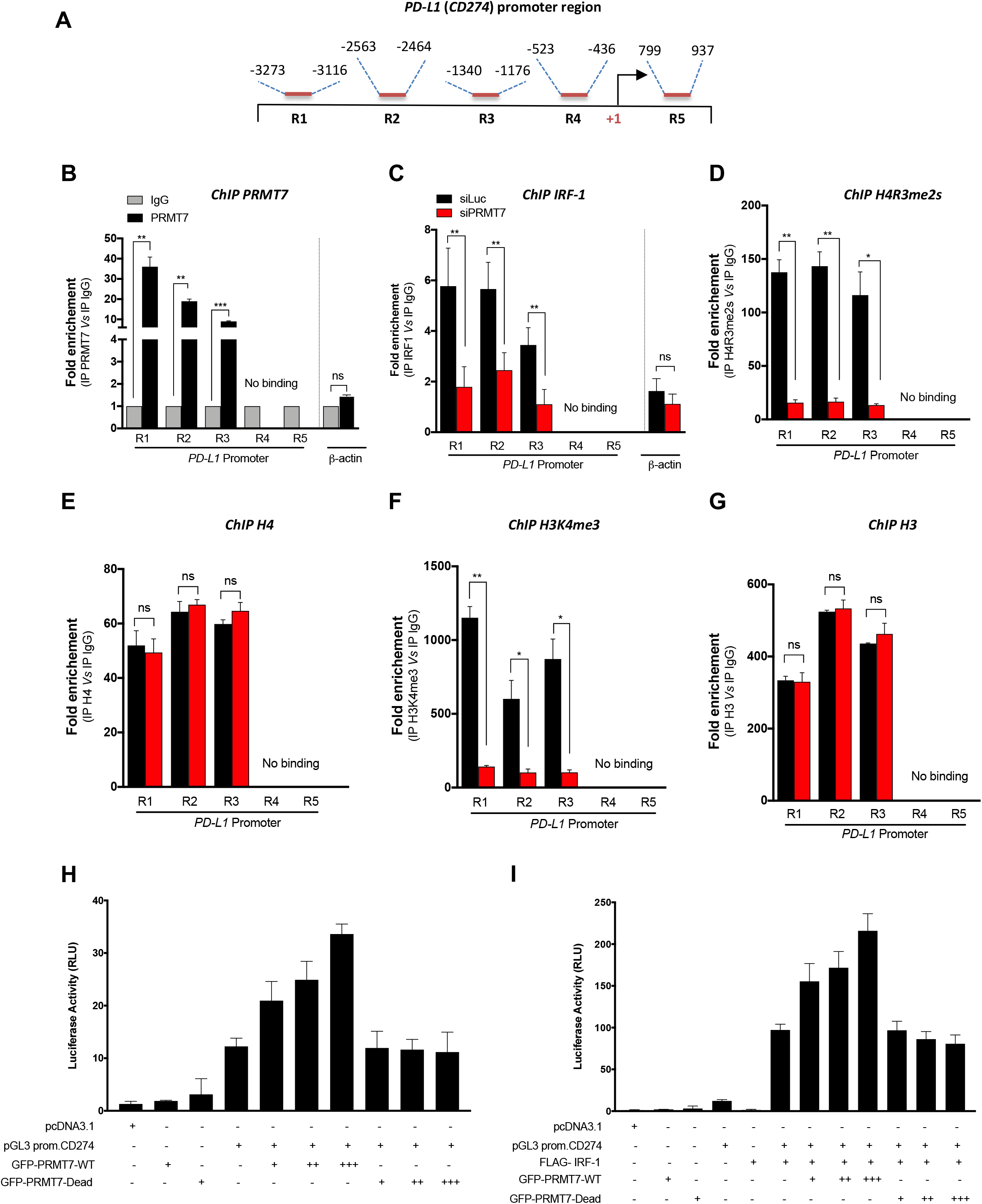
PRMT7 is an IRF-1 coactivator for the *PD-L1* promoter. **(A)** Schematic representation of the mouse *PD-L1* (*CD274*) promoter (−4,000 to +1,000 relative to *Cd274* transcription start site) showing the position of primers set used for different regions analyzed for ChIP (R1, R2, R3, R4 and R5). **(B)** Chromatin was prepared from B16.F10 melanoma cells and analyzed by ChIP with an PRMT7-specific antibody (red bars). The immunoprecipitated chromatin fragments were then analyzed by qPCR with primers spanning from −4,000 to +1,000 relative to the *PD-L1* promoter region. Results are represented as fold enrichment relative to IgG (black bars). **(C-G)** Analyses of distribution of IRF-1 **(C)**, H4R3me2s **(D)**, H4 **(E)**, H3K4me3 **(F)** and H3 **(G)** in the promoter regions of PD-L1 (R1, R2, R3, R4 and R5). siLuc and siPRMT7 B16.F10 melanoma cells are represented in black and red bars, respectively. Chromatin was immunoprecipitated using anti-IRF-1, anti-H4R3me2s and anti-H3K4me3 antibodies. Anti-H3 and anti-H4 were used as controls for Histone marks modifications and IgG isotype was used for mock precipitation to exclude non-specific enrichment. Subsequent qPCR was performed using promoter primer sets for PD-L1 and for β-actin (negative control). Data were first normalized to % of input, and the fold enrichment were then normalized to IgG. Asterisks denote significance in an unpaired *t* test (* *p* <0.05; \*\**p* <0.01; \*\*\**p* <0.001; *ns*: non-significant), and error bars denote SD. **(H)** HEK 293 **c**ells were transfected with pRLTK control plasmid (100 ng), pGL3 promoter CD274 luciferase reporter plasmid (200 ng) together with an increase amount of GFP tagged PRMT7-WT or PRMT7-Dead expression plasmids: 0 ng (−), 80 ng (+), 400 ng (++) and 2000 ng (+++). **(I)** HEK 293 cells were transfected with pRLTK control plasmid (100 ng), pGL3 promoter CD274 luciferase reporter plasmid (200 ng) and FLAG tagged IRF-1 expression plasmid (400 ng) together with an increase amount of GFP tagged PRMT7-WT or PRMT7-DEAD expression plasmids: 0 ng (−), 80 ng (+), 400 ng (++) and 2000 ng (+++). In all transfections, the pcDNA3.1 vector was added to bring the total plasmid to the same amount. Luciferase activity was analyzed 24h post-transfection using the Dual-Luciferase Reporter assay (Promega). Relative luciferase activity (RLU) was measured relative to the basal level of reporter gene in the presence of pcDNA3.1 vector after normalization with co-transfected RLU activity. Values are mean ± SD for three independent experiments.

Though established that PRMT7 can only form MMA, it has been shown to modulate the levels of H4R3me2s *in vivo* (Feng et al., 2013a; Blanc and Richard, 2017). Thus, we proceeded to examine the presence of H4R3me2s at the *PD-L1* promoter regions by ChIP analysis. Indeed, we detected a dramatic decrease of H4R3me2s in PRMT7-depleted B16.F10 cells regions R1-R3 (Fig. 4D), whereas the levels of histone H4 remained unchanged (Fig. 4E). Due to the ability of PRMT7 to influence neighboring histone marks, we assessed the presence of H3K4me3 (a hallmark of gene activation) at the *PD-L1* promoter. ChIP assay showed that H3K4me3 was decreased at the *PD-L1* promoter regions R1-R3 (Fig. 4F), but not total H3 levels (Fig. 4G). Our findings show that reduced H4R3me2s in siPRMT7 B16.F10 cells correlated with decreased H3K4me3 (activation mark) and *PD-L1* expression.

To functionally define whether PRMT7 is an IRF-1 coactivator for the *PD-L1* (also known as *CD274*) promoter, we transfected pGL3 promCD274 with IRF-1 and/or PRMT7 in HEK293 cells. The normalized luciferase activity of pGL3 promCD274 was increased up to 2-fold with augmenting amounts of wild type GFP-PRMT7-WT, but not enzyme inactive GFP-PRMT7 (GFP-PRMT7-Dead, Fig. 4H). Moreover, FLAG-IRF-1 increased the activity by ∼100-fold of the pGL3 promCD274 reporter gene (Fig. 4I), as expected (Garcia-Diaz et al., 2017). Importantly, augmenting amounts of the GFP-PRMT7 further increased luciferase activity (>200-fold), while the co-transfection of GFP-PRMT7-Dead had no effect (Fig. 4I). Our data show that PRMT7 has co-activator activity, as it potentiates the activity of IRF-1 at the *PD-L1* promoter.

### PRMT7 negatively regulates the IFN-γ pathway, antigen presentation and chemokine signaling

To identify other genes regulated by PRMT7, we performed a transcriptomic analysis of siLuc and siPRMT7 B16.F10 cells exposed or not to IFN-γ (Fig. 5A, Supplementary Fig. S6, S7 and Dataset S2). Geneset enrichment analysis (GSEA) revealed that genes related to the IFN-γ signaling pathway, antigen presentation, and chemokine signaling were significantly enriched in siPRMT7 cells (Fig. 5A, 5B; Dataset S2). Inspection of the list of genes from the RNA-seq data revealed an elevation in the IFN signaling response genes (*Trim25, Oas2, Trim21, Stat1, Nlrc5, Irf7*, and *Oas3*) in unstimulated siPRMT7 cells. Many of the IFN-γ responsive genes suppressed by PRMT7 were relevant to innate immunity and encoded key chemokines essential for recruitment of effector T cells. We also identified that several T cell attractant chemokines (*Cxcl1, Cxcl2, Ccl2, Ccl5* and *Ccl8*) were upregulated in the RNA-seq data of the siPRMT7 cells and these were confirmed by RT-qPCR (Fig. 5A, 5C-G).

**Figure 5:**
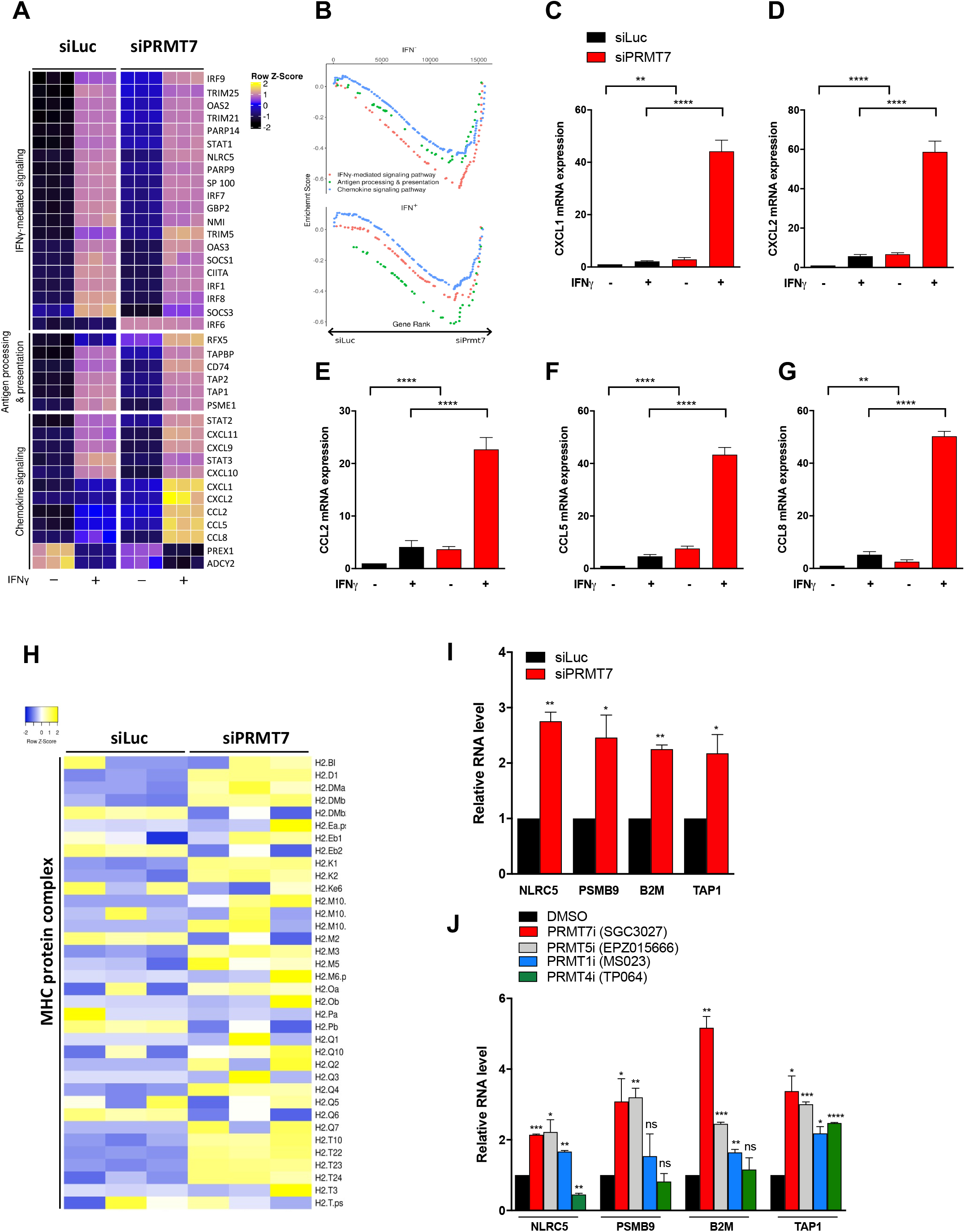
siPRMT7 B16 cells have up-regulated IFN pathway, antigen presentation and chemokine production. **(A)** siLuc and siPRMT7 B16.F10 cells (n=3 per group) were subjected to RNA-seq analysis. Heat map showing expression value (z-score based on cufflink count) of IFN genes, antigen processing and chemokine signaling genes with or without IFN-γ treatment (100 ng/ml) for 24h. **(B)** Gene set enrichment analysis of IFN-γ signaling pathway antigen processing and presentation and chemokine signaling pathway in siLuc and siPRMT7 cells. **(C-G)** RT-qPCR validation of genes identified from the RNA-seq dataset. Fold-change analysis using some selected genes: *Cxcl11***(C)***, Cxcl2* **(D)***, Ccl2* **(E)***, Ccl5* **(F)** and *Ccl8* **(G)** before and after IFN-γ treatment in siLuc (black) and siPRMT7 (red) B16.F10 cells. The fold-change in gene expression levels, before and after treatment, were calculated using the comparative cycle threshold (ΔΔCT) method and values were normalized to *Gapdh* mRNA levels as an internal control. Triplicates were used per biological sample. Bar graphs represent the mean fold-change ± SD. Statistical significance was calculated by unpaired student *t* test (\*\**p* <0.01; \*\*\*\**p* <0.0001). **(H)** Heat map showing expression value (z-score based on cufflink count) of all genes categorized in GO term ‘MHC protein complex’ in siLuc and siPRMT7 B16.F10 cells. **(I)** RT-qPCR analysis of genes implicated in antigen presentation (*Nlrc5, Psmb9, B2m and Tap1*) in siLuc and siPRMT7 B16.F10 cells. Bar graphs represent the mean fold-change ± SD. Data are representative of three independent experiments. Statistical significance was calculated by unpaired student *t* test (\**p* <0.1; \*\**p* <0.01). **(J)** RT-qPCR analysis of same transcripts analyzed in **(I)** in B16.F10 cells treated with the indicated PRMT inhibitors for 48h (SGC3027: 10 µM; EPZ015666: 5 µM; MS023: 600 nM and TP064: 3 µM). Bar graphs represent the mean fold-change ± SD. Data are representative of three independent experiments. Statistical significance was calculated by unpaired student *t* test (\**p* <0.1; \*\**p* <0.01; \*\*\**p* <0.001; \*\*\*\**p* <0.0001; *ns*: non-significant).

We also noted that in the siPRMT7 RNA-seq data, MHC-I coding genes, required for efficient display of antigens to effector T-lymphocytes (Dhatchinamoorthy et al., 2021), were upregulated in siPRMT7 cells (Fig. 5H). Moreover, known regulators of MHC class I genes, *Nlrc5* (nucleotide-binding oligomerization domain-like receptor family caspase recruitment domain containing 5) (Kobayashi and van den Elsen, 2012) and its target genes *Psmb9* (proteasome 20s subunit beta 9), *B2*m (Beta-2 microglobulin) and *Tap1* (antigen peptide transporter 1) were also upregulated in siPRMT7 or SGC3027 treated B16 cells (Fig. 5I, 5J). Interestingly, treatment of B16 cells with the PRMT5 inhibitor (EPZ015666) showed a significant increase in all MHC class I related genes (Fig. 5J), as described previously (Kim et al., 2020). A similar trend was observed with MS023 (PRMT1 inhibitor), except for *Psmb9* (Fig. 5J). In contrast, the CARM1 inhibitor (TP064) decreased *Nlrc5* expression, and increased *Tap1* mRNA levels, but did not affect *Psmb9* and *B2m* expression levels. These finding suggest that PRMT7 negatively regulates MHC-I gene expression, which then limits antigen presentation and enhances tumor evasion.

### Deletion of PRMT7 induces ‘viral mimicry’ through ERVs, dsRNA, and stress granule (SG) formation

Gene Ontology (GO) enrichment analysis of the differentially expressed genes between siLuc and siPRMT7 B16.F10 melanoma cells revealed top categories including 1) innate immune response, and 2) defense response to viruses (Supplementary Fig. S8). Many reports have linked endogenous retroviral elements (ERVs) to the activation of innate immune functions via IFN transcription and the regulation of tumor responses to host immunity (Chiappinelli et al., 2015; Roulois et al., 2015; Kassiotis and Stoye, 2016). To explore this possibility, we hypothesized that the IFN antiviral response induced in siPRMT7 cells might occur through the upregulation in the expression of ERVs and other retrotransposons in sense and antisense directions. First, we measured the levels of ERVs and IFN genes by RT-qPCR in sgCTL and both sgPRMT7-1 and sgPRMT7-2 B16.F10 melanoma cells. We observed in sgPRMT7 cells an upregulation of a number of ERV transcripts (ERVs: *MuERV-L, IAP, MuSD and Line-1*), IFN genes (IFNs: *Ifn-α, Ifn-β* and *Il-28*) and IFN-stimulated genes (ISGs: *Oasl, Isg15, Rig-1/Ddx58* and *Ifit*, Fig. 6A).

**Figure 6:**
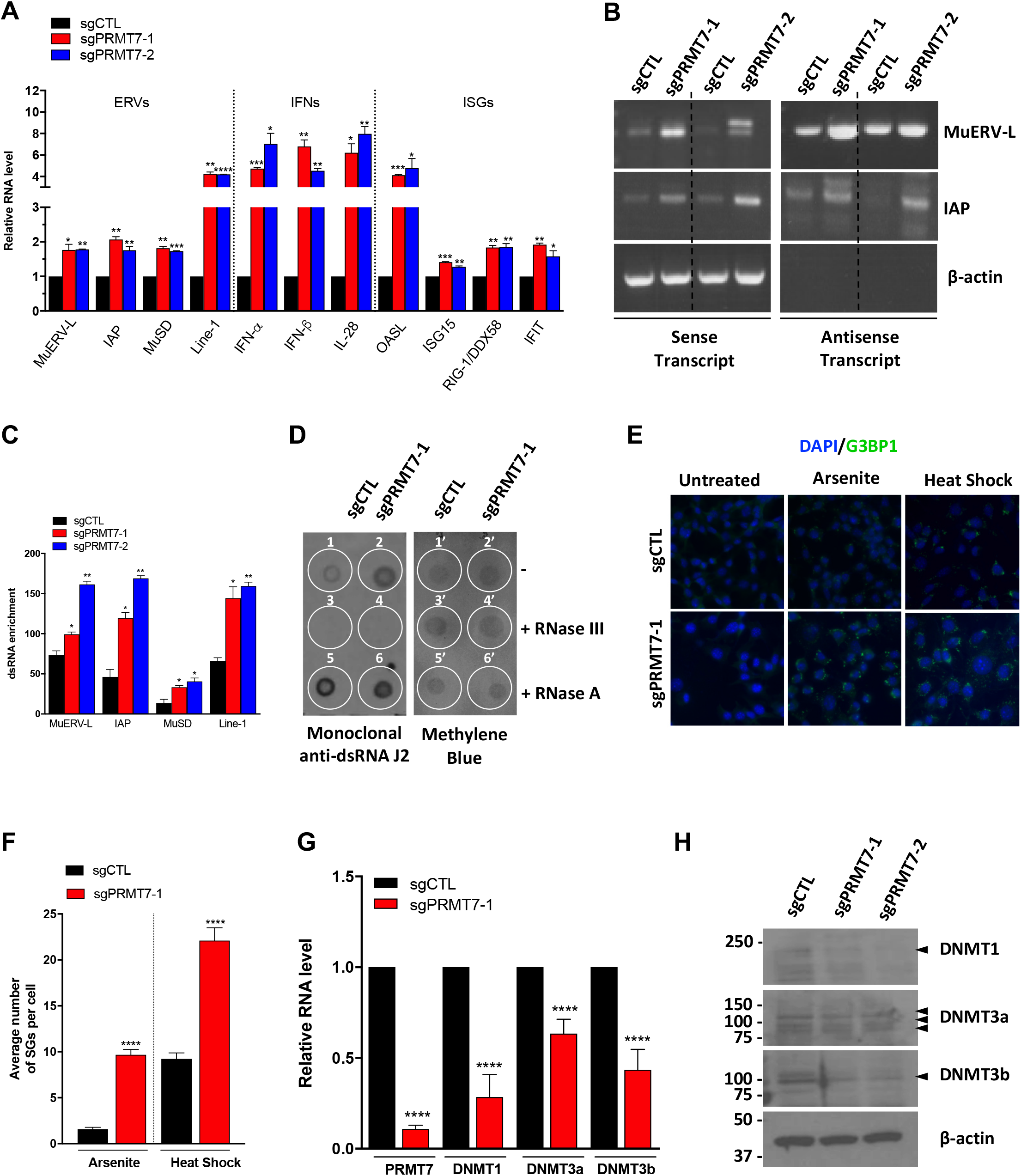
PRMT7 loss induces “viral mimicry” by regulating ERVs, dsRNA accumulation and stress granule (SG) formation. **(A)** RT-qPCR analysis of selected retrotransposons, IFNs and ISGs transcripts in sgCTL, sgPRMT7-1 and sgPRMT7-2 B16 cells. Bar graphs represent the mean fold-change ± SD. Data are representative of three independent experiments. Statistical significance was calculated by unpaired student *t* test (\**p* <0.1; \*\**p* <0.01; \*\*\**p* <0.001; \*\*\*\**p* <0.0001). **(B)** The assessment of both sense and antisense transcripts of selected ERVs (*MuERV-L* and *IAP*) using strand-specific primers for RT-PCR (TASA-TD technique) in sgCTL, sgPRMT7-1 and sgPRMT7-2 B16 cells. β-actin was used as a negative control for antisense transcription. A representative experiment is shown of three independent experiments. **(C)** dsRNA enrichment of *MuERV-L IAP, MuSD and Line-1* retrotransposons in sgCTL, sgPRMT7-1 and sgPRMT7-2 B16 cells by RT-qPCR analysis following RNase A treatment. **(D)** Total RNA extracted from sgCTL and sgPRMT7-1 B16 cells were treated with Mock, RNase III or RNase A (under high salt condition: 350 mM NaCl), dotted on Hybond N+ membrane and immunoblotted with the J2 antibody and visualized by methylene for loading control. Dots are denoted by numbers: 1, 3, 5 for sgCTL and 2, 4, 6 for sgPRMT7-1 cells nontreated (dots 1 and 2), treated with RNase III (dots 3 and 4) or RNase A (dots 5 and 6). **(E)** sgCTL and sgPRMT7-1 B16 cells were incubated with 0.5 mM sodium arsenite for 1h or 45°C (heat shock) treatment for 30 min. Cells were then fixed with 4% PFA and immunostained using anti-G3BP1 antibodies. A representative IF image is shown 60x magnification. DAPI, 4’,6-diamidino-2-phenylindole, was shown in blue as indicated. **(F)** The average number of SGs per cell of the staining done in **(E)** was quantified using image J software and presented as a bar plot (*n*=60 to 70 cells per condition). Bar graphs show mean intensity ± SEM. Statistical significance was calculated by unpaired student t test (\*\*\*\**p* <0.0001). **(G)** RT-qPCR analysis of *DNMT* mRNAs (*Dnmt1, Dnmt3a* and *Dnmt3b*) in sgCTL, sgPRMT7 B16 melanoma cells. Bar graphs represent the mean fold-change ± SD. Data are representative of three independent experiments. Statistical significance was calculated by unpaired student *t* test (\*\*\*\**p* <0.0001). **(H)** Immunoblot of DNMT proteins (DNMT1, DNMT3a and DNMT3b) in sgCTL, sgPRMT7-1 and sgPRMT7-2 B16 cells. β-actin was used as the loading control. A representative experiment is shown out of three independent experiments. The molecular mass markers are indicated in the left in kDa. The DNMT bands are shown with arrowheads.

We next examined whether bidirectional transcription producing sense and antisense transcripts of the murine subtype of ERVs: *MuERV-L* (Benit et al., 1997) and *IAP (intracisternal A-particles)* (Qin et al., 2010) could be detected using the TAG-aided sense and antisense transcript detection (TASA-TD) assay (Henke et al., 2015). Indeed, we detected higher levels of sense and antisense transcripts for *MuERV-L* and *IAP* in sgPRMT7 compared to sgCTL cells, but not *β-actin*, used as negative control (Fig. 6B), suggesting a role for PRMT7 in transcriptionally silencing ERVs. Such bidirectional expression are known to generate dsRNAs (Su et al., 2012) that trigger IFN responses (Gantier and Williams, 2007; Okamura and Lai, 2008; Berrens et al., 2017). To monitor if PRMT7 loss causes dsRNA accumulation, we treated total RNA from sgCTL, sgPRMT7-1 and sg-PRMT7-2 B16.F10 cells with RNase A, under high salt condition to cleave ssRNA and preserve the dsRNA. Indeed, RT-qPCR data showed dsRNA enrichment for a number of ERVs and other retrotransposons (*MuERV-L, IAP, MuSD, Line-1*) in sgPRMT7 cells compared to sgCTL (Fig. 6C). The dsRNA specific anti-J2 antibody detected dsRNA in an RNase III-dependent manner by RNA dot blot (Fig. 6D). The presence of intracellular dsRNAs in sgPRMT7 cells was higher than in sgCTL cells (compare dots 1 and 2, Fig. 6D). Together, these results demonstrate an accumulation of dsRNAs in the absence of PRMT7.

Sensing of the dsRNA, implicated in the innate immune response, was shown to be facilitated by stress granule (SG) formation (Burgess and Mohr, 2018). The later have antiviral activity and can mediate innate immunity through the SG nucleation component G3BP1 (DeWitte-Orr et al., 2009; McCormick and Khaperskyy, 2017). Thus, we investigated whether sgPRMT7 cells had increased G3BP1-positive SGs. Indeed, sgPRMT7 cells had increased number of SGs after sodium arsenite or 45°C heat shock treatment (Fig. 6E, 6F). Taken together, loss of PRMT7 in B16.F10 cells undergo a “viral mimicry” response with the upregulation of ERV expression, dsRNA, and increased presence of SGs.

### sgPRMT7 B16 cells have reduced DNMT1, DNMT3a, and DNMT3b expression

DNMT inhibitors are known to upregulate immune signaling through inducing ERVs in primary tumors, thus enhancing the sensitivity of tumors to immunotherapy (Chiappinelli et al., 2015; Zhang et al., 2017). As PRMT7 is known to influence DNA methylation (Jelinic et al., 2006), we wanted to investigate whether PRMT7 affected the expression of DNMTs in B16.F10 melanoma cells. Our transcriptomic analysis (RNA-seq) showed an ∼2-fold reduction in the expression of *Dnmt1* and a slight decrease in *Dnmt3a* and *Dnmt3b* expression (Dataset S1). RT-qPCR in sgCTL and sgPRMT7 B16 cells confirmed the reduced expression of *Dnmt1*, *3a* and *3b* mRNAs in sgPRMT7 cells (Fig. 6G). Moreover, the protein levels of DNMT1, 3a and 3b were also reduced in both sgPRMT7-1 and sgPRMT-2 compared to sgCTL cells (Fig. 6H). Taken together, our data suggest that PRMT7 regulates the expression of DNMTs and may indirectly affect DNA methylation and gene expression.

### Clinical relevance of PRMT7 expression in response to ICI therapy in human melanoma patients

To assess the clinical significance of our findings, we analyzed whether *PRMT7* mRNA expression can be used as a predictor of ICI response. RNA-seq data from two cohorts of melanoma patients treated with anti-PD-1 therapy were analyzed (Hugo et al., 2016; Riaz et al., 2017). Transcriptomic database of 28 patients treated with anti-PD-1 therapy showed that patients presenting a complete response to ICI treatment had a low mRNA level of *PRMT7* at pre-treatment (Hugo et al., 2016), suggesting lower *PRMT7* predicts a better ICI outcome (Fig. 7A, 7B). The same analysis was performed for a melanoma cohort treated with Nivolumab (anti-PD-1) (Riaz et al., 2017) and the data showed that low *PRMT7* expression level was correlated with a better response to ICI (Fig. 7C, D). The patients with a complete response (CR) showed the lowest level of *PRMT7* in pre-treatment biopsy and 29 days after the immunotherapy treatment, compared to other groups (PR: Poor Response; SD: Stable Disease; and PD: Progressive Disease). Interestingly *PRMT7* gene expression was more pronounced in patients with poor responses (Fig. 7C, 7D).

**Figure 7:**
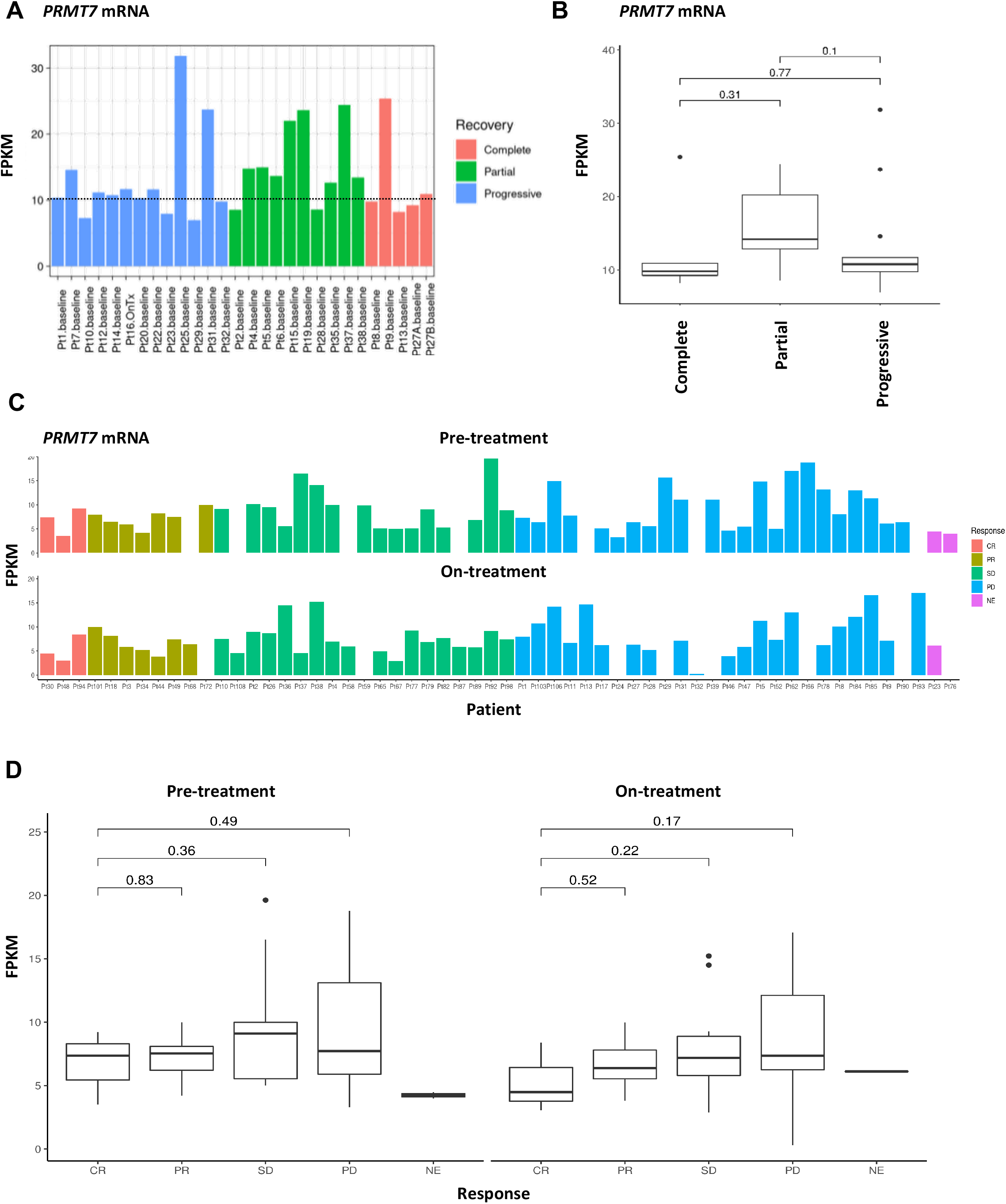
PRMT7 expression is inversely correlated with the response to ICI in human melanoma patients. **(A)** Plot of FPKM gene expression values showing the correlation between PRMT7 mRNA expression in patients treated with ICI therapy. n=28 cases grouped according to whether they receive complete (pink; n=5), partial (green; n=10) or progressive recovery (blue; n=13). The FPKM values were obtained from the GEO accession GSE78220. **(B)** Box plots showing the FPKM values for each case reported in (a). *p* values by Wilcoxon test are shown. **(C)** Plot of the FPKM gene expression values for PRMT7 showing the correlation between PRMT7 in patients before (upper panel) and during (lower panel) Nivolumab treatment. n=58 cases grouped according to whether they showed a complete response (CR, orange; n=3), partial response (PR, olive; n=8) or stable disease (SD, green; n=19), progressive disease (PD, blue; n=26). 2 patients were non evaluable (NE, pink; n=2). The FPKM values were obtained from the GEO accession GSE91061. **(D)** Box plots showing the FPKM values for each case reported in (**C**). *p* values by Wilcoxon test are shown.

In addition, we performed CD3 immunohistochemical (IHC) staining on formalin-fixed paraffin-embedded (FFPE) patient derived melanoma samples treated with anti-PD-1 plus carbotaxol (n=9) to establish an immune score. Our data showed that patients who responded better to ICI (GR: Good Responders) presented a higher immune score (2-moderate, 3-severe), compared to PR patients presenting a weak immune score (1-focal) (Table 1). These findings suggest a positive correlation between CD3^+^ T cells (immune infiltration) with the ICI outcome. Moreover, we performed an IHC staining against PRMT7 on same FFPE patient derived melanoma samples and found that in patients who exhibited a poor response (PR) to ICI therapy, 50% of melanomas stained positively for PRMT7 (Table 1). In contrast, in patients who were GR to ICI, their melanomas had a higher immune score (2-moderate, 3-severe) than PR, and only 20% of melanomas stained positive for PRMT7. Moreover, we showed that the expression of PRMT7 in human cancers negatively correlated with T cell cytotoxicity markers in TCGA datasets using TIMER (Tumor Immune Estimation Resource) method (Li et al., 2017) (Supplementary Fig. S9). We found that lower expression of *PRMT7* is correlated with higher cytotoxic activity contributed mainly by CD8^+^ T cells and granzyme B (GZMB) in skin cutaneous melanoma (SKCM) (Supplementary Fig. S9A, S9B). Also, we highlighted a global “partial” negative correlation between PRMT7 expression and the abundance of six subsets of immune infiltration cells including B cells, CD8^+^ T cells, CD4^+^ T cells, macrophages, neutrophils and dendritic cells in SKCM (Supplementary Fig. S9C), suggesting an important role for PRMT7 in reformatting the tumor immune microenvironment in melanoma. Taken together, our findings show a direct correlation between PRMT7 expression, immune infiltration and clinical response.

**Table 1:** Human melanoma correlative study. IHC staining towards CD3 and PRMT7 was established on nine human melanoma patient samples (FFPE tumor sections) treated with anti-PD-1 and carbotaxol. The level of CD3 and PRMT7 protein expression in the melanomas were scored and grouped according to whether they received clinical benefit from immunotherapy or not. 5 good responding tumors (GR: Good Responders) and 4 non-responding tumors (PR, Poor responders). The immune score was obtained from semi-quantitative prevalence of CD3^+^ cells noted as absent (0), focal (1), moderate (2) or severe (3). For the PRMT7 staining, the score was noted as POS for a positive PRMT7 nuclear staining or as NEG for a negative PRMT7 nuclear staining (absence of PRMT7 expression: low or undetectable).

| ***Patient ID*** | ***Response to ICI*** | ***PRMT7 Staining*** | ***Immune Score (CD3)*** |
| --- | --- | --- | --- |
| 01-003 | PR | POS | 1 |
| 01-013 | PR | NEG | x |
| 01-022 | PR | NEG | 2 |
| 02-023 | PR | POS | 1 |
| 01-002 | GR | NEG | 2 |
| 01-004 | GR | NEG | 3 |
| 02-012 | GR | POS | 3 |
| 02-015 | GR | NEG | 3 |
| 02-025 | GR | NEG | 2 |

## Discussion

In the present manuscript, we identify PRMT7 as a regulator of immunotherapy sensitivity for melanoma B16.F10 cells. CRISPR/Cas deletion of PRMT7 (sgPRMT) in B16.F10 cells resulted in enhanced adaptive immunity following anti-CTLA-4 and PD-1 treatment and smaller tumor formation when injected subcutaneously in syngeneic mice. The small tumors observed with sgPRMT7 had increased infiltration of CD8^+^ T cells with a decrease in G-MDSCs and M-MDSCs *in vivo*. Moreover, the sgPRMT7 generated melanomas had increased pigmentation associated with a melanocytic differentiated phenotype. Transcriptomic analysis by RNA-seq showed that PRMT7 is a regulator of gene expression for the IFN pathway, antigen presentation, and chemokine signaling. Additionally, we show that PRMT7 regulated the expression of PD-L1, DNMTs (1, 3a, 3b) and melanocytic markers MITF, Melan-A and GP100. Mechanistically, we show that PRMT7 serves as a coactivator for IRF-1, as PRMT7 was required for optimal IRF-1 chromatin recruitment, and the presence of the H4R3me2s and H3K4me3 histone activation marks on the *PD-L1* promoter. PRMT7 deficient cells had increased expression of transcripts derived from repetitive elements and resulting dsRNAs, mimicking a viral response. Finally, we show an inverse correlation between PRMT7 expression and ICI response in melanoma patients. These findings suggest that therapeutically inhibiting PRMT7 sensitizes melanoma to ICI therapy (Fig. 8, see model).

**Figure 8:**
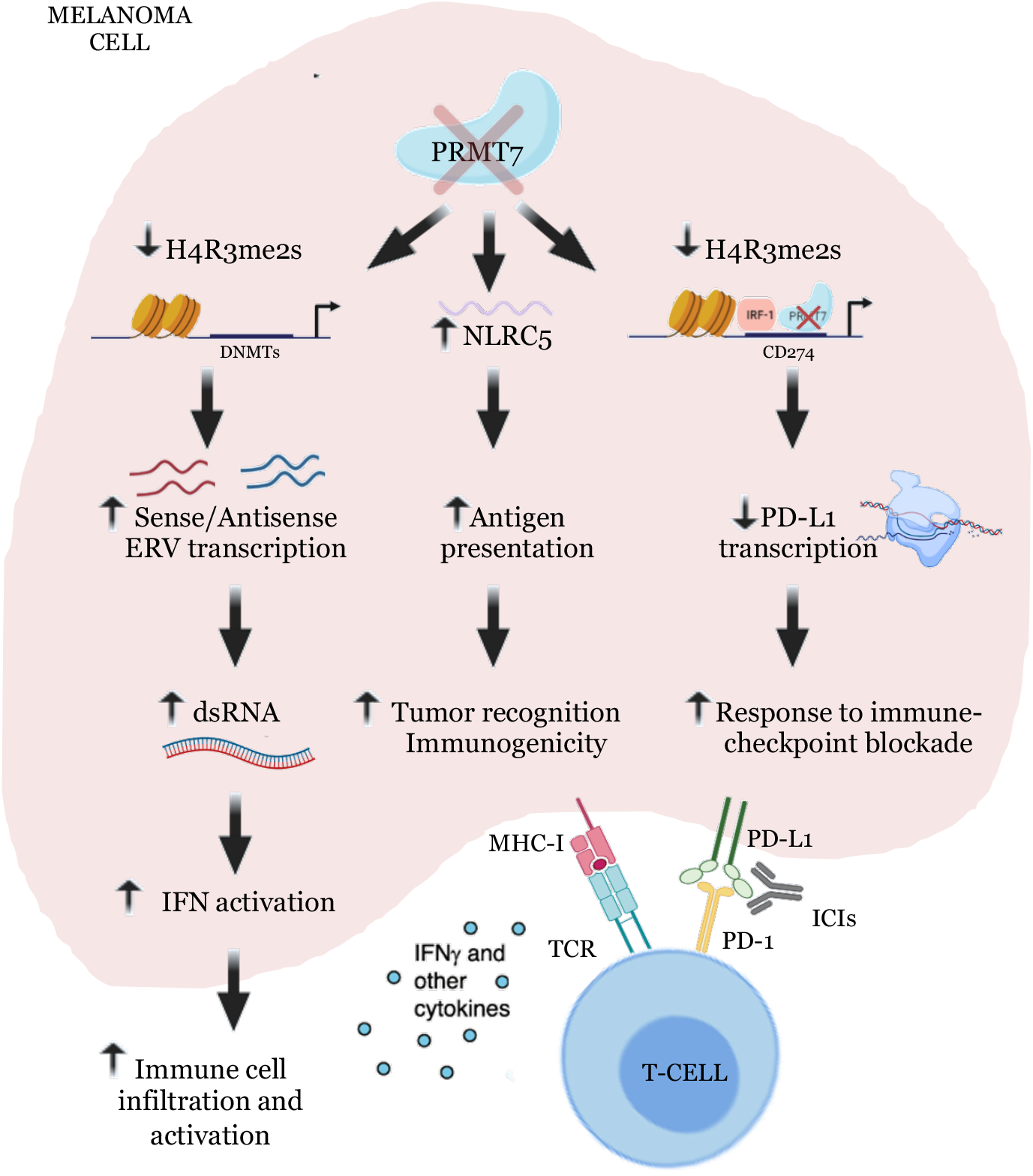
Proposed model for PRMT7 function in sensitizing melanoma to immunotherapy. PRMT7 deletion or inhibition in melanoma enhances tumor immunogenicity and sensitivity to cancer immunotherapy by stimulating ERVs, activating the IFN response, increasing antigen presentation and cytokine expression as well as decreasing *PD-L1* levels. This occurs through the decrease presence of H4R3me2s on *PD-L1* promoter and influencing DNA methylation.

Elevated expression of PD-L1 in cancer cells is a fundamental way to escape host immunity (Cha et al., 2019). Therapeutic antibodies designed to block the PD-1/PD-L1 interaction have emerged as one of the most efficient strategies to reverse this effect (Pardoll, 2012). In addition to this, epigenetic regulation such as histone acetylation (Woods et al., 2015; Hogg et al., 2017) and histone methylation (Lu et al., 2017; Toyokawa et al., 2019) play a crucial role in regulating PD-L1 expression. IRF-1 is a major transcription factor responsible for the constitutive and IFN-γ inducible PD-L1 expression (Garcia-Diaz et al., 2017) and IRF1-deficient tumor cells are unable to upregulate PD-L1 expression (Shao et al., 2019). Our findings that PRMT7 is an essential co-activator of IRF-1 to regulate the levels of H4R3me2s at the *PD-L1* promoter provide a new therapeutic mechanism for the regulation of PD-L1 expression. PRMT7-mediated H4R3me2s decrease at *DNMT3b* and *CDKN1a* promoters has been observed in muscle stem cells (Blanc et al., 2016). PRMT7 also influences the methylation of H4R3 levels at the *BCL6* gene and negatively regulates its expression for germinal center formation and plasma cell differentiation (Ying et al., 2015). The H3K4me3 decrease at the *PD-L1* promoter is consistent with PRMT7 regulation of mixed-lineage leukemia 4 (MLL4) catalyzed H3K4me3 level (Dhar et al., 2012).

Since PRMT7 expression levels allosterically influences the amount of PRMT5-catalyzed methylation of histone H4R3 (Jain et al., 2017), we first assumed that in PRMT7-deficient B16.F10 melanoma, H4R3me2s on the *PD-L1* promoter might be regulated via PRMT5. However, we observed that siPRMT5 B16.F10 cells showed an opposite response to PRMT7 deficient cells, with an increase in *PD-L1* mRNA expression, as recently reported (Kim et al., 2020). Furthermore, neither PRMT5, nor its cofactors, MEP-50, RIOK, COPR5, or pICIn were identified in the CRISPR/Cas screen for regulators of GVAX and PD-1, where PRMT7 was identified (Manguso et al., 2017). Taken together, these findings suggest that PRMT7 functions in a PRMT5-independent mechanism to regulate *PD-L1* expression.

We observed an elevated infiltration of CD8^+^ T cells and a decreased presence of G-MDSCs and M-MDSCs in sgPRMT7 melanomas *in vivo*. High levels of G-MDSCs and M-MDSCs are known to promote an immunosuppressive environment in skin cancer (Fujimura et al., 2012). Furthermore, IFN-γ is known to enhance cytotoxic T lymphocyte (CTL) function (Bhat et al., 2017) and inhibit MDSC function (Medina-Echeverz et al., 2014). A robust IFN-γ response in NSCLC patients and melanoma patients treated with ICIs is accompanied with a significantly longer progression-free survival (Higgs et al., 2018). Tumor cell loss of the *IFNGR1* gene results in resistance to anti-PD-1 (Shin et al., 2017) and anti-CTLA-4 therapies (Gao et al., 2016). PRMT5 deficiency (Kim et al., 2020; Ma et al., 2021), like our PRMT7 data, upregulate the IFN pathways, MHC-I related genes (*Nlr5, B2m, Bsmp9, Tap1*), and chemokine production. PRMT5 was shown to regulate the cGAS/STING pathway, known to limit the expression of IFN regulated genes and cGAS-mediated immune response (Kim et al., 2020; Ma et al., 2021). Thus, PRMT5 and PRMT7 regulate similar pathways i.e., IFN pathways, but using different strategies.

DNMT inhibitors increase the expression of ERVs in cancer cells to activate the innate antiviral response (Chiappinelli et al., 2015). We found that PRMT7 positively regulates DNMTs expression at the mRNA and protein levels, suggesting that PRMT7 loss in melanoma triggers ERVs through the expression of DNMTs. Actually, PRMT7 has been shown to influence the expression of DNMT3b in muscle stem cells (Blanc et al., 2016) and it is known to regulate DNA methylation (Jelinic et al., 2006). Furthermore, links between histone modifications and DNA methylation are known (Lehnertz et al., 2003; Esteve et al., 2006; Vire et al., 2006; Wu et al., 2021). The recruitment of DNMT3a to the H4R3me2s histone mark is known to facilitate DNA methylation and gene expression (Zhao et al., 2009).

In conclusion, our findings show that PRMT7 loss elicits anti-tumor immunity associated with increase immunogenicity and T cell infiltration. Since PRMT7 inhibitors and ICIs are in the same pathway, PRMT7 inhibitors may be effective in cases of ICI resistance. Our data provide the impetus for further drug development for more effective PRMT7 inhibitors (Szewczyk et al., 2020), as these can potentially be combined with immune-based therapies to achieve synergy. Future studies will be directed at ascertaining the use of PRMT7 inhibition across different cancer types and to examine if PRMT7 could be used as a biomarker for ICI responsiveness. Although our study focuses on PRMT7 inhibition; the possibility to target other type of PRMTs may enhance the efficiency of immunotherapy. Actually, a phase I clinical trial is ongoing to test PRMT5 inhibitors (GSK3326595) in combination with anti-PD1 (Wu et al., 2021).

## Acknowledgements

We would like to thank Dr. Xiaoru Chen for the expert technical assistance with the tumor slicing and Christophe Goncalves for helpful discussions and Dr. David Fisher (Harvard Medical School, Boston, Massachusetts) for his generous donation of the MITF antibody. This work was funded by a Canadian Institute of Health Research FDN-154303 awarded to S.R. N.S. was a recipient of a McGill fellowship (The Rosenberg/ Unger/ Schwarzbard/ Kallchman Hematology Research Award) and now holds a fellowship award from the Fonds de la recherche en santé du Québec (FRQS).

## Author contributions

N Srour: conceptualization, data curation, formal analysis, methodology, writing—original draft and project administration; O D Villarreal: conceptualization, resources, data curation, software, formal analysis, methodology, writing—review, and editing; Z Yu: supervision, methodology, writing—original draft and project administration; S Preston: resources, methodology; W H Miller, Jr.: resources; M M Szewczyk: resources, methodology; D Barsyte-Lovejoy: resources, methodology, project administration, writing—review, and editing; H Xu: conceptualization, resources and data curation; S V del Rincón: supervision, resources, methodology, project administration, and writing—original draft, review, and editing; S Richard: conceptualization, supervision, funding acquisition, project administration, and writing—original draft, review, and editing.

## List of Supplementary Materials

### Supplementary Figures

**Figure S1:** Loss-of-function screen identifies PRMT7; a target that increases the efficacy of immunotherapy.

**Figure S2:** PRMT7 expression in human cancer.

**Figure S3:** Cell viability determined by MTT assay after 24, 48 and 72 hours of incubation for sgCTL and both sgPRMT7-1 and sgPRMT7-2 B16.F10 cells *in vitro*.

**Figure S4:** PRMT7 inhibitor (SGC3027) reduces tumor volume *in vivo*.

**Figure S5:** Gating strategy for the immune composition of the tumor microenvironment in sgCTL control and sgPRMT7 B16.F10 cells *in vivo*.

**Figure S6:** RNA-seq data analysis in siLuc and siPRMT7 with or without IFN-γ treatment.

**Figure S7:** Changes in gene expression upon the specified treatment for each condition.

**Figure S8:** Gene Ontology analysis in siLuc and siPRMT7.

**Figure S9:** Expression of PRMT7 is negatively correlated with T cell cytotoxicity markers in TCGA datasets.

**Figure S10:** Uncropped Western blots.

**Dataset S1:** CRISPR/Cas9 results reported in Manguso et al. and re-analyzed by MoPAC software of three groups: Gvax+PD Vs TCRa, Gvax Vs TCRa and Gvax+PD Vs Gvax.

**Dataset S2:** RNA-sequencing data of PRMT7 proficient (siLuc) and deficient (siPRMT7) B16.F10 melanoma cells.

**Dataset S3:** List of antibodies and primers used.

### Materials and Methods

#### *In vivo* CRISPR screening analysis in B16.F10 tumor cells

The differential analysis of the CRISPR screen performed by Manguso et al. (Manguso et al., 2017) was carried out with the MoPAC v3.1 (Modular Pipeline for Analysis of CRISPR screens) R package (Gao et al., 2019). In brief, the log-fold-change at both the sgRNA and gene levels were first obtained from the table of read counts with a quality control module. Afterwards, a normalization module was used to compute unbiased measures of sgRNA and gene essentiality. The statistical significance was assessed based on: (1) the Z-score of the differential gene essentiality, (2) Student’s *t*-test applied to the biological replicates of gene essentiality and (3) a novel bidirectional version of MAGeCK’s αRRA algorithm (Li et al., 2014)applied to the differential sgRNA essentiality. The MoPAC tool is publicly available at https://sourceforge.net/projects/mopac/.

#### Generation of sgPRMT7 B16.F10 cells

The B16.F10 melanoma cell line was kindly provided by Dr. Michael Pollak (McGill University). These cells were subjected to CRISPR/Cas9-Mediated knockout of PRMT7 by transient co-transfection of the Cas-9 single guide RNA (sgRNA)-GFP plasmid (Addgene # Px458) and the PRMT7 sgRNA plasmid (IDT: # 270436658), targeting the exon four with the following gRNA sequence: 5’-*AAA ATA CTA CCA GGG TAT CCG GG*-3’. 5×10^5^ cells were plated in a six-well plate and were co-transfected the following day using 2µg of pX459 (Cas-9) plasmid DNA and 2µg of PRMT7 sgRNA plasmid DNA using Lipofectamine 3000 (Invitrogen) according to manufacturer’s instruction. 24 hours later, GFP positive cells were isolated using FACS-ARIA sorter (Beckton Dickinson). After selection, cells were grown for 14 days *in vitro* before being tested for the deletion of PRMT7 by immunoblotting and subsequently the deletion junction sequenced by Sanger DNA Sequencing. Two positive clones (sgPRMT7-1, sgPRMT7-2) and one negative clone (sgCTL) were selected to be used for the experiments.

#### Cell lines

B16.F10 melanoma cell lines were maintained in Dulbecco’s modifies Eagle’s medium (HyClone), supplemented with 10% Fetal Bovine Serum (FBS: HyClone), 1% penicillin/streptomycin (Multicell) and 1% sodium pyruvate (Multicell) in a 5% CO_2_ incubator at 37 °C. Cells were routinely tested for mycoplasma contamination.

#### Animals

All mouse procedures were performed in accordance with McGill University guidelines, which are set by the Canadian Council on Animal Care. Seven to twelve-week-old wild type female C57BL/6J mice were obtained from Jackson laboratories (Stock No: 000664). A colony of B6.129S2-Tcra^tm1Mom^/J (Tcra^−/−^) T cell-deficient mice were also obtained from Jackson laboratories (Stock No: 002116). Mice were age-matched to be 7 to12 weeks old at the time of tumor inoculation. Mice were subcutaneously injected with 1×10^6^ cells/100µl into the right flank on day 0. On day 3, 6, 9 and 12, mice were treated with 100µl of monoclonal anti-PD-1 (anti-mouse CD279, clone: RMP1-14, Cat #BE0146, *InVivo*MAb) and 100µl of anti-CTLA-4 (anti-mouse CD152, clone: 9H10, Cat #BE0131, *InVivo*MAb) *via* intraperitoneal injection. Tumors were measured every two to three days beginning on day 3 after challenge until the time of sacrifice. Measurements were taken manually with a caliper by collecting the longest dimension (length) and the longest perpendicular dimension (width). We estimated the tumor volume with the formula: (L×W^2^)/2. CO_2_ inhalation was used to euthanize mice 15 days after tumor inoculation for tumor collection. For the PRMT7 inhibitor injection *in vivo*: 7 to12 weeks old mice were subcutaneously injected with 1×10^6^ cells/100µl B16.F10 melanoma cells into the right flank on day 0 and then intratumorally injected with 10µM of DMSO, SGC3027N (control compound) or SGC3027 (PRMT7 inhibitor) on day 7, 8, 9 and 10. Tumor size was measured 96 hours after the last drug injection and calculated as described above.

#### Cell culture, transfections and treatments

Melanoma cell lines were seeded into six-well plates on day 1, targeting 70–80% of confluence on the day of analysis. On day 2 after siRNA transfection, cells were exposed to 100 IU/ml interferon gamma (MACS Miltenyi Biotec #130-105-785) for 24 hours. For siRNA and vector transfections, B16.F10 cells were transfected using Lipofectamine RNAiMAX (Invitrogen) and Lipofectamine 3000 (Invitrogen) respectively, according to the manufacturer’s instruction. All siRNAs (20 to 40 nM) were purchased from Dharmacon and the sequences are as follow: siPRMT7 (siGenome SMARTpool mouse PRMT7#214572 siRNA, Catalog ID: M-053294); *siPRMT7#1: 5’-GGA CAG AAG GCC UUG GUU C-3’; siPRMT7#2: 5’-GAG CGG AGC AGG UGU UUA C-3’; siPRMT7#3: 5’-UCA GCU AUG UUG UGG AGU U-3’; siPRMT7#4: 5’-GUA GCU UCC UAU AGA CUG A-3’*; siPD-L1 (siGenome SMARTpool mouse CD274#60533 siRNA, Catalog ID: M-040760); *siPD-L1#1: 5’-GAU AUU UGC UGG CAU UAUA-3’; siPD-L1#2: 5’-GAG GUA AUC UGG ACA AAC A-3’; siPD-L1#3: 5’-GAG CCU CGC UGC CAA AGG A-3’; siPD-L1#4: 5’-GAA UCA CGC UGA AAG UCA A-3’* and siPRMT5 (siGenome SMARTpool mouse PRMT5#27374 siRNA, Catalog ID: M-042281); *siPRMT5#1: 5’-CAA CCG AGA UCC UAU GAU U-3’; siPRMT5#2: 5’-GGA AUA CGC UAA UUG UGG G-3’; siPRMT5#3: 5’-GUC CGU GCC UGU CGG GAA A - 3’; siPRMT5#4: 5’-CAG UUU AUC AUC ACG GGA A-3’*. The *siRNA 5’-CGU ACG CGG AAU ACU UCG AdTdT-3’*, targeting the firefly luciferase (GL2) was used as control (siLuc).

#### RT-qPCR

Total RNA from cells were isolated with TRIzol (Invitrogen) according to manufacturer’s instruction. After digestion with DNase I (Promega), 1 μg of total RNA was converted to cDNAs using M-MLV reverse transcriptase (Promega). Real-time quantitative PCRs were performed using PowerUp SYBR Mastermix (Life Technologies #A25742) on 7500 Fast Real-Time PCR System (Applied Biosystem). Results were normalized as described in the figure legends using the ΔΔct method. Primers used in this study are outlined in Dataset S3.

#### Protein extracts and immunoblot analysis

Whole lysates from B16.F10 melanoma cells were prepared in 2x Laemmli buffer and boiled at 100°C. Equal amounts of protein samples were loaded, separated on 10% SDS-PAGE gels, transferred to nitrocellulose membranes using an immunoblot TurboTransfer system (Bio-Rad) and probed with corresponding antibodies listed in Dataset S3. Immunoblot signals were detected using chemiluminescence (Perkin Elmer).

#### Flow cytometry analysis

B16.F10 cells were transfected with the corresponding siRNA and then treated or not with 100IU/mM interferon gamma (Mouse IFN-γ, Cat#130-105-785, MACS, Miltenyibiotec) for 24 hours. On day 3, cells were blocked with Fc-Block and thein stained with anti-PD-L1 antibody (CD274-clone: #558091, BD-Pharmingen). For immune cell composition analysis: Primary tumors were collected on day 15, weighed, mechanically diced, incubated with collagenase P (2 mg/ml, Sigma-Aldrich) and DNase I (50 μg/ml, Sigma-Aldrich) for 10 min, and pipetted into a single-cell suspension. Cells were then blocked with anti-mouse CD16/32 antibody (BioLegend) and stained with indicated antibodies (Dataset S3) as well as a Live/Dead discrimination dye (BD Biosciences). Data were subsequently acquired at the LSR Fortessa flow cytometer and results were analyzed using FlowJo software.

#### Immunofluorescence (IF)

B16.F10 tumors were fixed for 24 hours in 10% neutral-buffered formalin and then permeabilized in 70% ethanol overnight. Briefly, tissue sections were blocked in 10% normal goat serum in PBS with 0.3% Triton X-100/PBS solution for 1 hour, followed by primary antibody incubation overnight at 4°C. For G3BP1 staining, B16 cells were growing on glass coverslips and treated with 0.5 mM sodium arsenite (NaAS2O3, Sigma S1400) for 1 h or heat shock at 45°C for 30 min. Cells were then fixed for 10 min with 4% paraformaldehyde (PFA), washed with PBS and permeabilized for 5 min with 0.25% Triton X-100 in PBS. Coverslips were then incubated with blocking buffer containing 5% FBS for 1h, and incubated with G3BP1 antibodies for 2h at RT. After three washes, slides and coverslips were incubated with corresponding fluorescent secondary antibodies for 1 hour at RT and mounted with IMMU-MOUNT (Thermo Scientific) mounting medium containing 1µg/ml of 4′,6-diamidino-2-phenylindole (DAPI). Images were taken using a Zeiss M1 fluorescence microscope and analyzed by ImageJ.

#### Immunohistochemical Staining (IHC) and scoring

Human melanoma patient samples were obtained from the Sir Mortimer B Davis Jewish General Hospital, Montreal, Quebec, Canada. Patients with melanoma were treated with a combination of anti-PD-1 and carbotaxol and pre-treatment tumor tissues were obtained for IHC staining performed on a Ventana Discovery Benchmark XT. Briefly, formalin-fixed, paraffin-embedded tumor sections were stained with CD3 (Ventana Benchmark: clone 2GV6 at 1:50) (Taube et al., 2014) and PRMT7 (Sigma #HPA044241 at 1:10) antibodies, followed by a standard Fast Red detection protocol. Hematoxylin-counterstained slides were mounted with coverslips. Staining intensity was determined by a clinically certified pathologist who was blinded to all clinical data and antibodies used for IHC.

#### Chromatin Immunoprecipitation (ChIP)

ChIP was performed as previously described (Mersaoui et al., 2019).using the SimpleChip plus Chromatin IP Kit (CST; Magnetic beads 9005) according to the manufacturer’s instruction. Briefly, formaldehyde cross-linked chromatin was prepared from 2×10^7^ B16.F10 melanoma cells, and the samples were immunoprecipitated with the corresponding antibodies (Dataset S3) overnight at 4°C and rabbit-IgG isotype control was used for mock precipitation to exclude any non-specific enrichment. The primers used for qPCR are listed in Dataset S3.

#### Dual-luciferase reporter assay

HEK293T cells were seeded in 24-well plates and then transiently co-transfected in triplicate with the pGL3-PD-L1 promoter-luc reporter (Plasmid #107003, Addgene) (Coelho et al., 2017) alone or with the indicated expression vectors (FLAG-IRF-1, GFP-PRMT7-WT or GFP-PRMT7-Dead). The FLAG tagged IRF-1 plasmid was a gift from Dr. Rongtuan Lin (Lady Davis Institute, McGill University). The GFP-tagged PRMT7-WT and the catalytically inactive mutant PRMT7-Dead plasmids were a gift from Dr. Mark T Bedford (University of Texas M.D. Anderson Cancer Center). The Renilla pRL-TK plasmid was used as an internal control and the total amounts of DNA were kept constant by supplementation with an empty vector (pcDNA3.1). After 24 h, cell lysates were harvested, and the relative luciferase units (RLUs) were measured using the dual-luciferase assay according to the manufacturer’s instructions (Promega #E1910). RLUs from firefly luciferase signal were normalized by RLUs from Renilla signal.

#### Strand-specific PCR for detection of sense and antisense ERV transcripts

The strand-specific PCR method was adapted from (Henke et al., 2015) (TASA-TD) and performed with the MultiScribe RT-PCR (Applied biosystems # 4366596) with some modifications. Briefly, the gene and strand specific primers (GSP) were synthesized with an extra TAG sequence at the 5’ end, which does not exist in the mouse genome (listed in Dataset S3). The first strand cDNA synthesis reaction was performed following these steps. 1 µg total RNA in 6 µl H2O was mixed with 1 µM TAG-GSP, 0.5 mM dNTP, 40 U RNase inhibitor, 100 U MultiScribe RT and 240 ng Actinomycin D (Sigma-Aldrich, #A9415) to a total volume of 20 µl; incubated at 42°C for 30 min and terminated at 85°C for 5 min. The resulting single sense or antisense cDNA/RNA hybrids were then treated with 2 U of recombinant RNase H (NEB #M0297S) to generate single strand cDNAs, followed by ethanol precipitation for cDNA purification. To amplify sense cDNA: a TAG primer and GSP sense (PCR) were used and to amplify antisense strand: a TAG primer and GSP antisense (PCR) were used. Sense and antisense specific PCRs for β-actin were used as an internal control (no antisense transcripts). The amplicons were visualized on 1.5% agarose gels.

#### dsRNA analysis by RT-qPCR

5 µg of total RNA extracted from B16.F10 cells was dissolved in 46 µl H2O and digested with 1U RNase A (Ambion #AM2270) under high salt condition: 3.5 µl NaCl (5 M stock) to a total volume of 50 µl and mixed well, followed by incubation for 30 min at 37°C. H_2_O was used as mock. Then, 1 ml TRIzol was added to the mixture to terminate digestion, followed by RNA extraction. The transcript expression of selected retrotransposons was measured by RT-qPCR with GAPDH as an internal control. The dsRNA-fold enrichment was calculated as the ratio of retrotransposon/GAPDH_RNaseA_/retrotransposon/GAPDH_mock_.

#### dsRNA analysis by J2 immunoblotting

Total RNA extracted from B16.F10 cells was digested with mock (H_2_O), RNaseIII (Thermo Fisher Scientific, #AM2290) according to the manufacturer’s instructions, or with RNaseA (Ambion, #AM2270) under high salt condition (350 mM NaCl) as described previously (Sheng et al., 2018). Briefly, equal volumes of purified and treated RNA were dotted on Hybond N+ membrane (GE Healthcare, #RPN303B), dried and auto crosslinked in a UV machine (Bio-Rad GS Gene linker) using the following program: 125mJoule/cm^2^ at 254 nM. The membrane was probed with J2 antibody at 4 °C overnight and ECL was applied for film development. For the loading control, membrane was stained for 30 min with methylene blue solution (0.3% w/v methylene blue + 30% v/v ethanol + 70% v/v H_2_O).

#### RNA sequencing and data analysis

RNA samples were purified using GenElute™ Mammalian Total RNA Miniprep Kit (RTN70, Sigma Aldrich). Total RNA was assessed for quality using an Agilent Tapestation 4200, and RNA sequencing libraries were generated using TruSeq Stranded mRNA Sample Prep Kit with TruSeq Unique Dual Indexes (Illumina, Hiseq4000, SR75 platform located at San Diego, CA; UCSD IGM Genomics Facility, La Jolla, CA). Samples were processed following manufacturer’s instructions, starting with 50 ng of RNA and modifying RNA shear time to 5 min. Resulting libraries were multiplexed and sequenced with 100 base pair (bp) to a depth of approximately 30 million reads per sample. Samples were demultiplexed using bcl2fastq v2.20 Conversion Software (Illumina, San Diego, CA). Reads were mapped to the Genome Reference Consortium Mouse Build 38 patch release 6 (mm10/GRCm38.p6: primary assembly) (Frankish et al., 2019) using STAR v2.4 (Dobin et al., 2013).

#### Gene expression analysis

Expression levels were estimated using HOMER V4.10 (Heinz et al., 2010). Afterwards, we employed DESeq2 (Love et al., 2014) to normalize the raw counts as rlog variance stabilized values, as well as to perform the differential expression analysis as previously described (Darbelli et al., 2017). For the volcano plot, genes were considered differentially expressed if they had an adjusted *p* value <0.05, a base mean higher than 100 and an absolute fold-change greater than 2. For the heat map, genes were considered differentially expressed if the samples with the highest and lowest expression are more than 2-fold different and one of the samples has 25 normalized reads (as in the HOMER tutorial).

#### Gene ontology

GO term enrichment analysis of the differentially expressed genes was performed through one or more of the following: (1) STRING v11.0 (Szklarczyk et al., 2019), (2) GSEA V3.0 (Mootha et al., 2003), (3) Enrichr, (4) DAVID, (5) IPA. The list of differentially expressed genes was compared to a background of expressed genes, consisting of all expressed genes in the complete dataset (defined as all genes with the DESeq2 base mean higher than the first expression quartile). For differentially expressed genes, upregulated and downregulated genes with a base mean higher than 100 and an absolute fold-change greater than 2 were used for the analysis. Publicly available gene expression data was obtained from the CCLE (Cancer Cell Line Encyclopedia and Genomics of Drug Sensitivity in Cancer, 2015).and/or TCGA (Barretina et al., 2012).

#### Quantification and statistical analysis

All experiments were repeated at least two to three times, except as specified otherwise. All data are presented as mean ± standard error of the mean (SEM). Graph Pad Prism Version 6 was used to generate plots and additional statistical analysis. Significance of comparison between two groups was assessed either by the unpaired or paired Student*-t test*. The use of the specific tests as well as the number of animals and experimental replicates has been reported in each figure legend. Statistically significant results were defined as follows: * *p* <0.05; \*\**p* <0.01; \*\*\**p* <0.001; \*\*\*\**p* <0.0001. Statistical analysis for RNA-seq was performed with DESseq2 for gene expression.

#### Data and software availability

The RNA-seq data is available at NCBI under accession number GSE157141.

